# Auditory-Vestibulomotor Temporal Processing and Crossmodal Plasticity for Musical Rhythm in the Early Blind

**DOI:** 10.1101/2020.03.23.987727

**Authors:** Jessica Phillips-Silver, John W. VanMeter, Josef P. Rauschecker

## Abstract

The auditory dorsal stream (ADS) is a cortical brain network responsible for sensorimotor control and integration, including spatiotemporal processing. Although spatiotemporal movement of the head and body involves input from the vestibular system, and despite the wealth of evidence for the strong coupling between the vestibular and visual systems, very little is known about how vestibular information is integrated with auditory-motor inputs in the ADS. There is also no evidence addressing to what extent auditory-vestibulomotor integration is affected by early visual deprivation. Using functional magnetic resonance imaging and motion capture technology we show that in a task of sensorimotor temporal processing (‘feeling the beat’), the ADS includes an extension to parietoinsular vestibular cortex (PIVC) and to subcortical regions including basal ganglia and vestibular cerebellum. This circuit is engaged after sensorimotor synchronization training, during beat recognition, and is preserved in the early blind. The strength of activation of PIVC in the early blind correlates with a measure of lifetime physical spatial activity, suggesting that experience with vestibular stimulation via physical spatial activities might compensate for any negative effects of early blindness, and thus reinforcing the potential beneficial effects of mobility training. Finally, rhythmic entrainment provides an effective tool for studying auditory-vestibulomotor integration and music appreciation, and for developing music- and movement-based interventions for early blind individuals.

---

The auditory dorsal stream (ADS) is known to be responsible for spatiotemporal anticipation and action (Rauschecker and Scott, 2009). To accomplish this, the ADS takes into account not only auditory information, but also the movement of the head and body. Thus, a main function of the ADS consists of the conversion of sensorimotor sequences into a unified experience in real time (Rauschecker, 2018). The processing of auditory spatial (“where”) and temporal (“when”) information that relies on the ADS is generally preserved, and can even be enhanced as a result of early visual deprivation in animals and humans (Rauschecker and Korte, 1993; Renier et al. 2010). For example, the early blind (EB) have preserved auditory spatial abilities (Renier et al. 2010; Collignon et al., 2011). They also show enhanced temporal processing in the form of auditory beat asynchrony detection—i.e., knowing when one beat in a rhythmic sequence is ‘out of time’ (Lerens et al. 2014). The crossmodal plasticity that is evident in early blindness, in which visual cortex is recruited during tasks involving auditory or tactile spatial processing, might help explain certain preserved or enhanced abilities in those modalities (Korte and Rauschecker, 1993; Sadato et al. 1996; Gougoux et al. 2005; Collignon et al. 2009; Dormal et al. 2012; Renier et al. 2014). However, much less is known about the role of the ADS or of crossmodal plasticity for temporal aspects of sensorimotor processing in the EB.

In any task involving the perception of spatiotemporal information as the head or body moves in space, vestibular information is critical. Seemungal and colleagues (2007) reported that early blindness resulted in deficient vestibular perception (velocity storage) and impaired performance on a spatial reorientation task (path completion) which requires an inferential spatial strategy that utilizes vestibular information. While Seemungal et al. found impaired overall ability in the EB group to utilize spatial mechanisms during vestibular navigation, the performance of two out of six blind individuals reached the level of sighted controls (SC), which corresponded with notably higher degrees of lifetime physical spatial activity as compared to their blind peers. Physical spatial activities referred to activities of daily living that have a vestibular component, such as participation in sports. To assess the level of lifetime physical activity and compute a score for each blind individual, Seemungal et al. used a questionnaire that inquired about the amount of time spent engaged in physical activities, the types of activities and their estimated degree of vestibular stimulation (e.g., degree of whole-body reorientation), and the individuals’ own perceived level of ability as compared with their blind peers. Each of these variables was assigned a point value, and from this the researchers computed a score of lifetime physical spatial activity for each participant. The authors suggested that while early visual deprivation may tend to inhibit vestibular stimulation and perception (for example, through reduced movement), extensive physical spatial activity might compensate for this and even lead to superior vestibular navigation in the EB (Seemungal et al. 2007). Engagement in physical spatial activity is thus likely to be not only a matter of quality of life, but a potential factor in determining levels of enhancement or deficiency in sensorimotor spatiotemporal processing in the EB.

A model for examining the perception of spatiotemporal information as well as the sensorimotor integration required for producing a precisely timed action is musical beat processing—also known as rhythmic entrainment (Phillips-Silver and Keller, 2012). In rhythmic entrainment, individuals perceive the beat of a rhythmic sequence and synchronize their body movement in time (Phillips-Silver et al. 2010, 2011). This phenomenon requires the functions of predictive timing and motor planning that are characteristic of the ADS (Zatorre et al. 2007; Rauschecker and Scott, 2009; Grahn et al., 2011; Grahn, 2012; Kung et al., 2013; Patel and Iversen, 2014). Beat-based rhythmic sequences are known to recruit a ‘motor-cortico-basal-ganglia—thalamo-cortical’ (mCBGT) circuit for timing, beat processing and action simulation in non-human primates (Mendoza and Merchant, 2014; Merchant et al. 2014). These two circuits— ADS and mCBGT—describe essentially the same brain network for timing, with key roles for premotor cortex, supplementary motor area (SMA) and pre-SMA, as well as extensions to task-relevant cortical and subcortical regions such as auditory cortex, basal ganglia and cerebellum.

An area that has not yet been examined in investigations of the ADS or of the mCBGT circuit, however, is vestibular cortex, which is due to the fact that it is difficult to assess the effects of body motion inside an MRI scanner. Yet behavioral evidence of rhythmic entrainment in infants and adults indicates the integration of auditory, motor and vestibular signals (Phillips-Silver and Trainor, 2005; 2008; Trainor et al. 2009). The vestibular cortex thus warrants consideration as an extension of the ADS timing network in light of three points: 1) the ADS is essential for sensorimotor integration during movement of the whole body in space and time, 2) vestibular information is particularly important for perception during action (such as during music and dance) but has largely been neglected, and 3) early visual deprivation may have multifaceted effects on vestibular perception and function.

In the present view of rhythmic entrainment, if the integration of auditory, vestibular and motor inputs is sufficient for beat processing, then visual experience may not be required and thus early blind individuals should be able to achieve body synchronization and subsequent beat recognition. On the other hand, if rhythmic entrainment requires input from the visual system, then early visual deprivation should prohibit synchronization and beat recognition.

We used functional magnetic resonance imaging (fMRI) and motion capture technology in early blind (EB) and matched sighted control (SC) participants, to map the ADS timing network in a task of sensorimotor rhythmic entrainment. The present study is based on a behavioral paradigm in which participants are trained to listen to a repeating bistable (ambiguous) percussion rhythm, while bouncing continuously on every second (binary) or every third (ternary) beat of the rhythm. The movement training (which is recorded with a portable motion capture device and measured for synchronization) requires subjects to recognize test versions of the rhythm pattern that have binary or ternary acoustic accents, respectively (Phillips-Silver and Trainor, 2005, 2007, 2008). This paradigm has been used to show that full body movement influences the perception of the bistable auditory rhythm during training and causes infant and adult subjects to “feel the beat” of the ambiguous rhythm without having had acoustic accents. Here we implemented the same behavioral training outside of the scanner, so that participants learned via their body movement to interpret the bistable rhythm as having strong beats in either binary or ternary form. During this training we used a portable motion capture device—the Nintendo Wii remote—to record the participants’ acceleration data and analyze it for their synchronization as defined by the proportion of synchronized power (see Phillips-Silver et al. 2011). Immediately following the training, participants entered the scanner and performed the following tasks. First, they continued to listen to the bistable rhythm pattern while tapping along to the strong beat by pressing the button box (this was to provide sufficient practice and to have a task that included a motor response). Second, they heard test versions of the rhythm pattern that had binary or ternary accents and were asked to identify the test pattern that matched what they had heard during the training (this was the main experimental task, in which we would observe the neural circuit responsible for beat recognition). We predicted that the prior experience of rhythmic entrainment, by moving the body while learning to interpret the ambiguous rhythm, would subsequently activate the ADS circuit in EB and SC participants during the task of beat recognition. We predicted furthermore that the early blind would show evidence of crossmodal cortical plasticity by recruiting visual cortical areas for these musical tasks.

## Materials and Methods

### Participants

Fifteen early blind individuals (**Table 1**) and fifteen matched sighted controls were tested in the Laboratory of Integrative Neuroscience and Cognition at Georgetown University Medical Center in Washington, D.C. The EB participants (eleven females, four males) had a mean age of 39.4 years (range 18-63.9 years). Twelve were blind from birth; one was blind by the age of 1 year; one was blind at birth, gained some low vision and then lost it fully by age 2; and one had extremely low vision (only some light and form perception) at birth and was totally blind by the age of 8 years. All EB participants were blind as a result of peripheral damage [genetic eye disease (N=7), glaucoma (N=3), retinal disease (N=4) and corneal disease (N=1)]. They were totally blind (N=8) or did not have more than rudimentary sensitivity for light, form or motion (N=7). All EB participants read braille. The sighted control participants (mean age 40.3 years, range 19-65 years) were individually matched for age, sex and level of education. All participants gave written informed consent for the study, which was overseen by the Institutional Review Board of Georgetown University Medical Center.

**Table 1.** Demographic Information for Early Blind (EB) Participants, including music experience (in years) and Physical Spatial Activity (PSA) score.

| Participant | Move | Sex | Age | Education | Cause of Blindness | Age of Onset | Music (yrs) | PSA |
| --- | --- | --- | --- | --- | --- | --- | --- | --- |
| EB 1 | T | M | 38.1 | Graduate (masters) | Glaucoma | Birth | 15 | 9 |
| EB 2 | B | F | 41 | Graduate (masters) | Albanism, diabetes (genetic) | Birth | 11 | 8 |
| EB 3 | T | M | 46.1 | Bachelors | Retina detachments | Birth | 30 | 7 |
| EB 4 | B | M | 18.1 | HS | Leber Congenital Amaurosis (genetic) | Birth | 8 | 6 |
| EB 5 | T | F | 31.7 | HS (some college) | Glaucoma + cataracts | Birth, gained low vision, lost by 2 | 17 | 5 |
| EB 6 | B | F | 63.9 | Bachelors | Retinitis pigmentosa (genetic) | 1 year | 14 | 3 |
| EB 7 | T | F | 35.4 | Graduate (masters) | Cataracts (genetic) | Birth | 11 | 8 |
| EB 8 | B | F | 36.4 | Graduate (law) | Corneal disease (sclera cornea) | Birth | 8 | 8 |
| EB 9 | T | F | 32 | Graduate (PhD) | Leber Congenital Amaurosis (genetic) | Birth | 8 | 2 |
| EB 10 | B | F | 61.8 | Bachelors | Retinopathy of prematurity | Birth | 2 | 4 |
| EB 11 | T | F | 30.3 | Graduate (law) | Retinal blastoma cancer (genetic) | Birth | 7 | 8 |
| EB 12 | B | F | 25.9 | Graduate (law) | Retinopathy (optic nerve fibroplasia) | Birth | 21 | 4 |
| EB 13 | T | F | 32.2 | Graduate (masters) | Retina detachments, glaucoma | Nearly blind at birth; blind by 8 | <1 | 4 |
| EB 14 | B | F | 44.6 | HS (some college) | Glaucoma juvenile | Birth | 2 | 7 |
| EB 15 | T | M | 54.5 | Bachelors | Retinoblastoma | Birth | 30 | 5 |

Based on a questionnaire on lifetime experience with musical and physical spatial activities, we computed a physical spatial activity score for each participant (Seemungal et al. 2007). This scoring system divided the lifetime activities into three epochs, from 0-10 years, 11-18 years, and above 18 years of age, and each activity was assigned one point per epoch of regular occurrence. Participants’ descriptions of their own musical and physical abilities were assigned points as follows: below average (0 points), average (1 point), above average (2 points), exceptional (3 points). Finally, each type of activity was assigned points according to the extent to which the activity required whole-body spatial reorientation: Seemungal and colleagues termed these “vestibulospatial activities”. We included in this category activities such as downhill skiing, swimming and triathlons (3 points each), versus activities in which the body is relatively static but requires coordination of the hands or limbs, such as blind yoga, and hand bells (1 point each). The sum of points for the duration of activity (epochs), self-described ability level, and degree of vestibulospatial orientation required, made up the Physical Spatial Activity (PSA) score in **Table 1**. Number of years of music lessons or practice is listed in the table for each participant.

### Stimuli

Auditory rhythm stimuli were produced on a digital audio workstation (Cakewalk Sonar X3 Producer (Cakewalk, Inc., Boston, MA). Instrument timbres were created within Sonar using third-party drum kit and Latin percussion sounds from virtual instruments XLN Audio Addictive Drums 2 (XLN Audio, Stockholm, Sweden) and Toontrack Latin Percussion EZX, respectively (Toontrack, Umeå, Sweden). Stimuli were normalized for loudness by using the “Radio” function in batch normalization software WaveGain (John Edwards developer, open access), which adjusts volume levels based on RMS values, and then by manually fine-tuning individual stimulus levels. All stimuli were encoded in WAV audio file format. Auditory test stimuli were presented in the scanner using Sensimetrics S14 insert earphones (Sensimetrics Corporation, Gloucester, MA) with Sperian Thunder T1 26-decibel noise reduction headphones (Honeywell, Inc., Smithfield, RI).

### Training Stimulus

The training rhythm pattern was a bistable composite rhythm created by superimposing binary and ternary strong beat patterns (strong beats every second versus every third beat, respectively) of a six-beat polyrhythm sequence, in accordance with Phillips-Silver and Trainor (2005, 2007, 2008). The bistable rhythm was made by presenting a 6-beat background (at a tempo of 206 beats per minute) that consisted of a downbeat (snare drum timbre) presented every 1750 ms, and a microbeat (slapstick timbre) occurring every 292 ms (3.42Hz). Superimposed over this background was the ambiguous training rhythm, consisting of four snare drum tones with stimulus onset asynchronies of 583-292-292-583 ms. The training stimulus was presented on a loop for 69 repetitions of the rhythm, for a duration of 2 minutes.

### Beat Recognition Task Stimuli

The stimulus pairs for the Beat Recognition Task consisted of two versions of the training pattern, each of which contained acoustically accented beats (i.e., they were no longer ambiguous). One version had accents on every second beat—binary grouping, which corresponded to a 6/8 time signature with three metrical strong beats (i.e., BEAT-rest-BEAT-beat-BEAT-rest) occurring at 1.72Hz (103 beats per minute). The other version had accents on every third beat—ternary grouping, which corresponded to a 3/4 time signature with two metrical strong beats (i.e., BEAT-rest-beat-BEAT-beat-rest) occurring at 1.14Hz (68 beats per minute). This effect was achieved by keeping the accented tones (the ‘strong beats’) at the same intensity level as the snare drum tones in the training stimulus, while reducing the relative intensity level of the unaccented tones (the ‘weak beats’). This stimulus pair construction was applied across a variety of percussive timbres (e.g., conga, floor tom, hand claps, cowbell) and combinations, resulting in a diverse stimulus set in order to prevent habituation. Timbres varied between trials, but on each trial both stimuli in the pair (binary and ternary) were presented in identical timbres (for example, floor tom + cowbell), and participants were instructed to recognize the familiar pattern based on their training experience. Since the ambiguous training stimulus had been presented in snare drum + slapstick timbres only, the test stimuli required generalization to new timbres in order to promote challenge and engagement in the task of recognizing the familiar beat.

### Experimental Design and Statistical Analysis

#### Training

Before entering the scanner, participants were exposed to a brief (two minute) training period during which they were instructed to listen to and remember the ambiguous training stimulus, while moving along with the experimenter. Participants stood facing the experimenter with their hands placed on hers and followed as she bent her knees and bounced to a beat: half of the participants were bounced to the binary beat (1.72Hz), and half to the ternary beat (1.14Hz). We recorded the participants’ body motion with a portable motion capture device (the Nintendo Wii remote, which measures acceleration of body movement) in order to determine that they had bounced at the correct frequency according to the binary or ternary grouping (Phillips-Silver et al. 2011).

After the familiarization with the training pattern and movement, participants entered the scanner and performed the following tasks. The first task was Tapping, in which the participants listened to the ambiguous stimulus and tapped along by pressing on the button box. The main purpose of this task was to observe brain activation while listening to the ambiguous rhythm pattern and performing the motor action of tapping along according to their prior training. The second and main experimental task was Beat Recognition, in which the participants listened (without tapping) to pairs of test stimuli which were no longer ambiguous but had accents in binary versus ternary form, and identified which stimulus in the pair corresponded to the pattern they had learned during training. Each task was presented using a block design with alternating active and rest blocks of 15s each (**Figure 1**). There were 12 blocks per run for two runs, lasting 6 minutes each (total time per task, 12 minutes).

**Figure 1.**
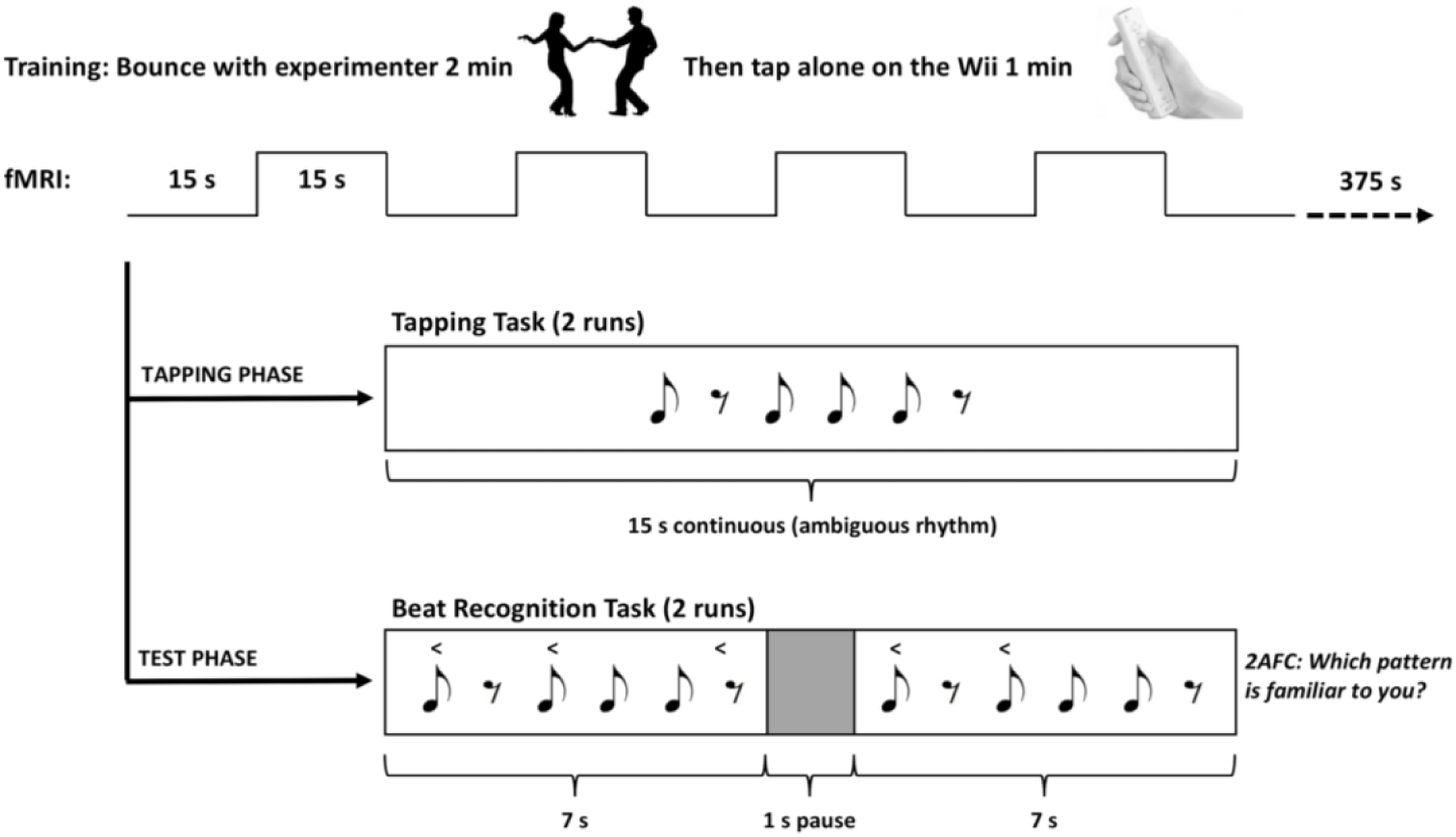
Behavioral and fMRI paradigm.

### Motion Capture Acquisition and Analysis

As in Phillips-Silver et al. (2011), body movement data were captured with an accelerometer contained in the remote control of the Nintendo wii, and recorded to a computer via bluetooth using the custom software Wii Data Capture (Version 2.2, University of Jyväskylä, Finland). The wii remote was strapped to the trunk of the participant’s body during the behavioral training and measured the acceleration of bouncing with a temporal resolution of 100 frames per second (10 ms), from which the beat-by-beat period of the vertical movement was computed. We defined the ‘bounce beat’ as the zero-crossings of the downward vertical acceleration (Toiviainen, Luck, and Thompson, 2010). We then applied a Fourier analysis to the continuous movement data, which enabled us to measure the overall proportion of synchronized power at the beat frequency of the auditory stimulus. This ‘proportion of synchronized power’ was the measure that determined the level of synchronization of body movement for each participant.

### Functional and 3D MRI Acquisition

#### MRI acquisition

Imaging data were acquired at the Center for Functional and Molecular Imaging at Georgetown University with an echo-planar imaging (EPI) sequence on a 3-Tesla Siemens Tim Trio scanner with a 12-channel head coil (flip angle = 90°, TR = 3 s, TE = 60 ms, 64 × 64 matrix). The FOV was 192 × 192 mm^2^ at the 64 × 64 matrix size; slice thickness was 2.8 mm with a 0.2 mm gap resulting in an effective voxel resolution of 3 mm^3^. Each volume of 47 slices was acquired using continuous sampling. A three-dimensional T1-weighted structural MPRAGE image (resolution 1 × 1 x 1 mm^3^) was acquired for each subject (TR = 2530 ms, TE = 3.5 ms, TI = 1100 ms, flip angle = 7**°**, field of view = 256 × 256, slice thickness = 1 mm, matrix size = 256 × 256).

### Image Analysis

Data analysis was performed using SPM version 12 for MatLab (Mathworks, Inc., Natick, MA). The fMRI data were preprocessed with slice timing correction and head motion correction. The computed statistical maps were overlaid on 3-D T1-weighted anatomical scans. Functional and anatomical data were co-registered and translated into MNI space using the Segment function. The fMRI data were spatially smoothed using a 6 mm^3^ gaussian filter. First-level statistical maps were computed for each task by convolving the canonical hemodynamic response function (HRF) with the task timing. Individual whole-brain contrast images were created per task. Second-level group comparisons were computed using an ANOVA. Images were family-wise error (FWE)-corrected for multiple comparisons, thresholded at p < 0.05, with clusters defined by an uncorrected threshold of .001 and an extent threshold of 10 voxels. The computed statistical maps were overlaid on 3-D T1-weighted anatomical scans.

## Results

### Motion Capture and Behavioral Task Performance

Results of the pre-fMRI motion capture showed that both SC and EB groups had significant proportions of synchronized power at the beat frequencies corresponding to the type of movement (Binary, Ternary) as led by the experimenter (Phillips-Silver et al. 2011). The proportion of synchronized power for each group was 0.84 (SD = 0.13) in the EB, and 0.86 (SD = 0.09) in the SC.

We first examined the fMRI data for the task Beat Recognition versus Rest, in order to observe the entire circuit that was engaged during this rhythmic entrainment task. In the Beat Recognition task, both SC and EB groups with Binary Movement training identified as familiar the test stimulus that had acoustic accents on every second beat, corresponding to the way they had bounced during training and tapped in the scanner to the ambiguous rhythm pattern (SC mean accuracy = 77.4%, SE = 9.8; EB mean accuracy = 78.6%, SE = 9.1; **Figure 2**). The SC group with Ternary Movement training also identified as familiar the test stimulus that corresponded to the way they had bounced during training and tapped in the scanner, with acoustic accents on every third beat (mean accuracy = 83.9%, SE = 9.1). However, results of the EB group with Ternary Movement training were much more variable—with a subset of the group identifying as familiar the Binary rhythm stimulus despite their training. Nevertheless, no differences in accuracy between Binary and Ternary Movement training groups, or between SC and EB groups, reached significance. A mixed ANOVA with factors Group (EB, SC) and Movement Type (Binary, Ternary) as independent variables and accuracy on the Beat Recognition task as dependent variable revealed no significant effect of Group (*F*_(1,23)_ = 2.45, *p* > 0.05) or Movement Type (*F*_(1,23)_ = 0.98, *p* > 0.05), and no significant Group by Movement Type interaction (*F*_(1,23)_ = 3.8, *p* > 0.05).

**Figure 2.**
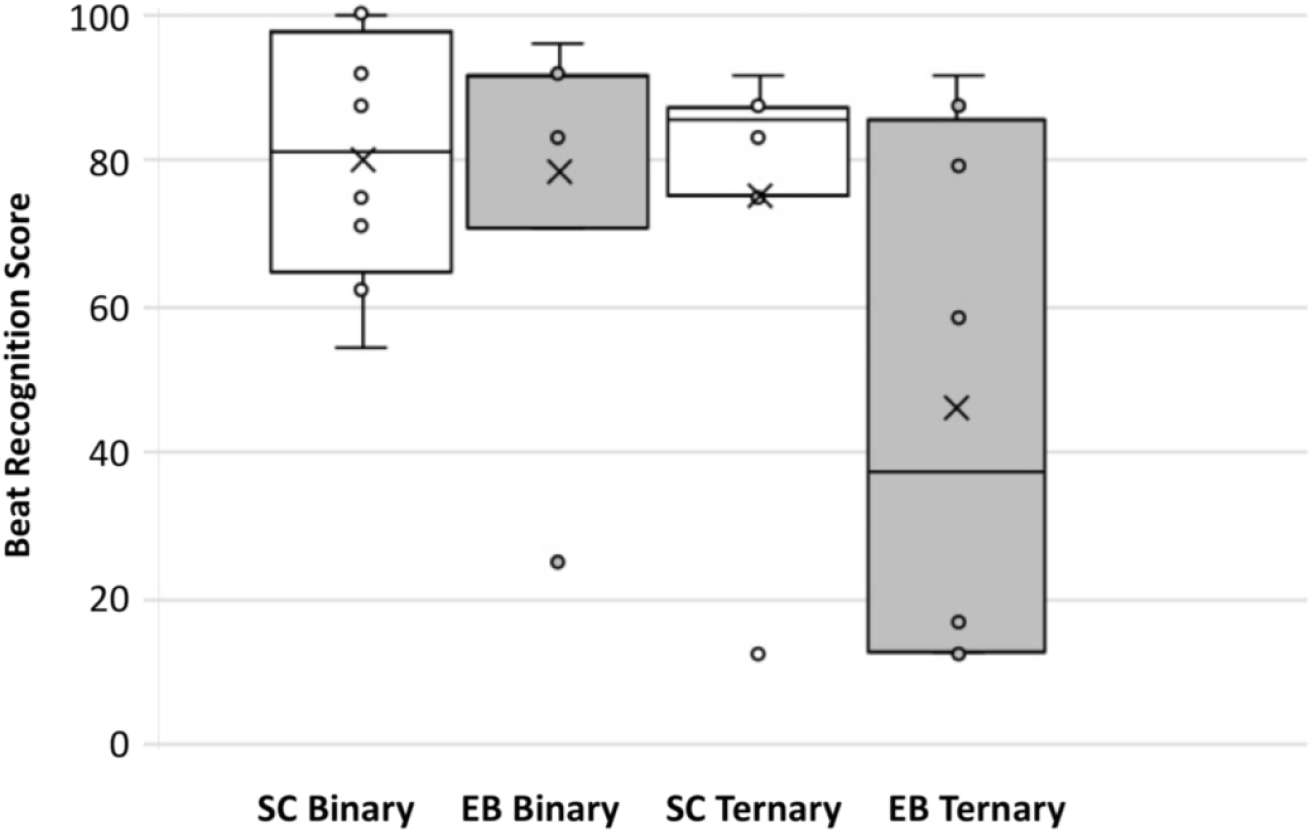
Performance on Beat Recognition shown as proportion of stimulus choice (binary, ternary) determined by movement type (binary versus ternary bounce, tap) in EB and SC.

### fMRI Results

#### Crossmodal Plasticity in the Blind: Sensorimotor Temporal Processing Activates Visual Cortex

In the EB group, the sensorimotor temporal processing task recruited visual cortex. The contrast EB > SC, whole-brain corrected at *p*_(FWE)_ < 0.05 (**Table 2, Figure 3**, left panel) showed cross-modal activation for both Beat Recognition and Tapping tasks, in right middle occipital gyrus (BA19), and right cuneus (BA19). This was the only significant difference found in brain activations between the EB and the SC groups, and thus the subsequent analyses were performed on EB and SC groups combined.

**Table 2.**
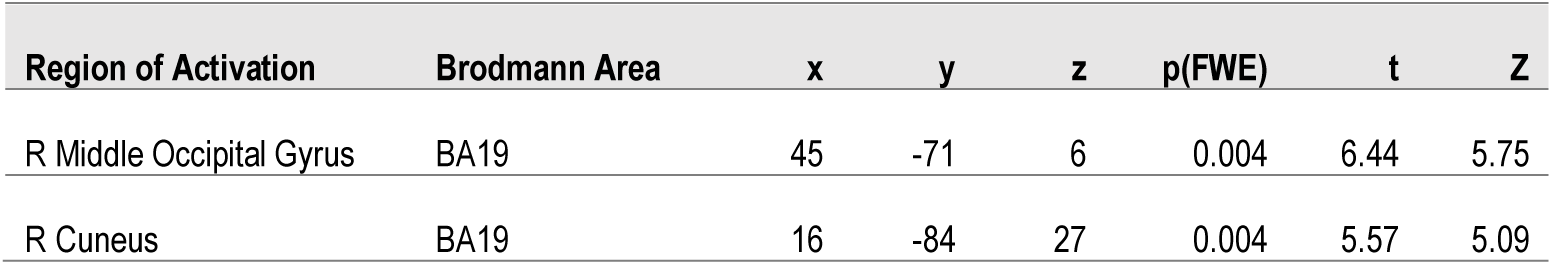
Talairach coordinates for brain regions active in EB > SC, whole-brain corrected at *p(FWE)* < .05.

**Figure 3.**
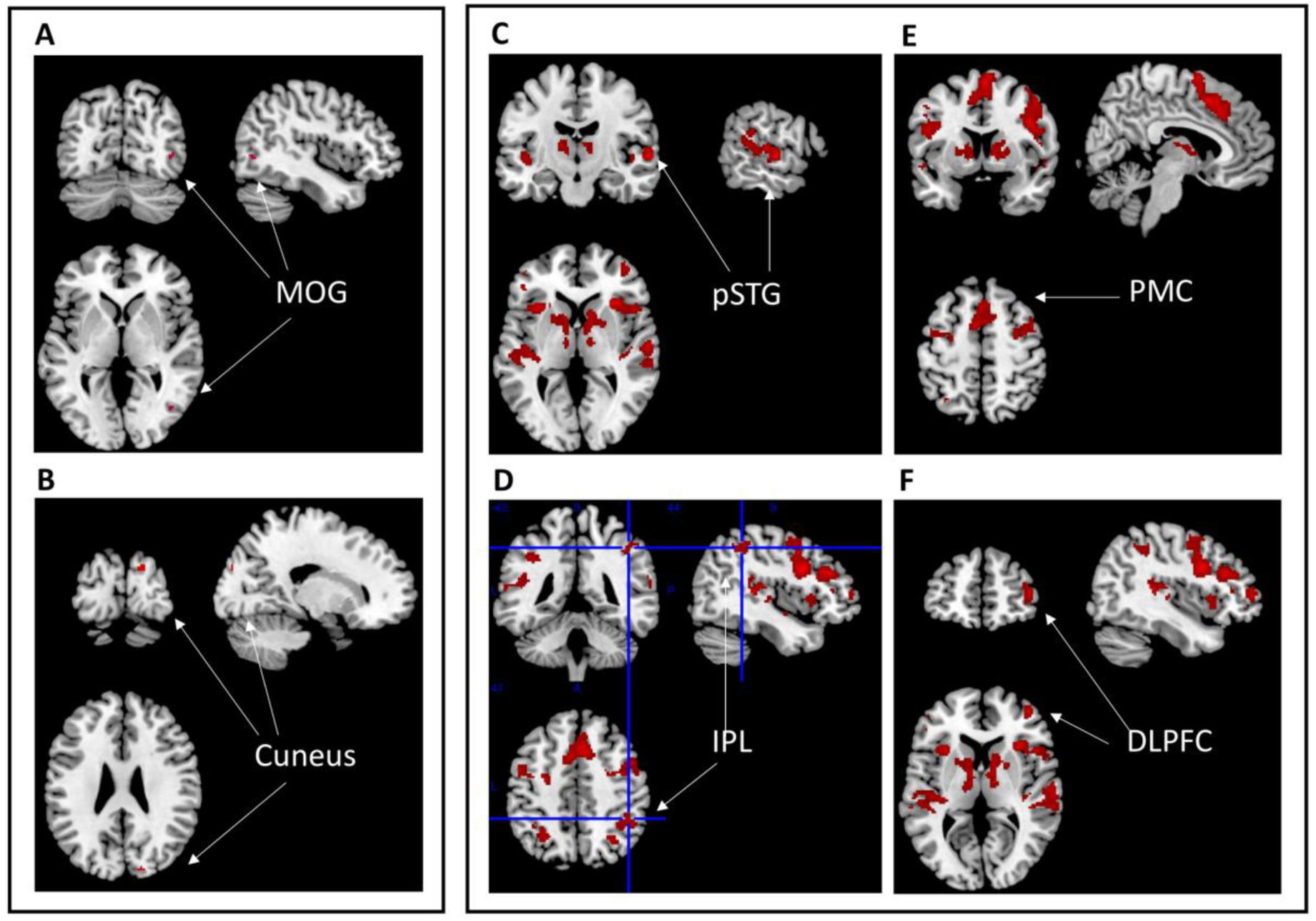
**(A-B)** Crossmodal plasticity for sensorimotor temporal processing in EB. The contrast EB > SC, whole-brain corrected at *p*_(FWE)_ < 0.05, showed cross-modal activation across Beat Recognition and Tapping tasks in right middle occipital gyrus (BA19), and right cuneus (BA19). **(C-F)** Beat Recognition activated the ADS in EB and SC. The contrast Beat > Rest, whole-brain corrected at *p*_(FWE)_ < 0.01, activates bilateral pSTG (BA22), bilateral IPL (BA40), bilateral premotor cortex (BA6/8), and right DLPFC. Also activated were related subcortical regions of the auditory-motor timing circuit: bilateral basal ganglia (claustrum), left anterior and bilateral posterior cerebellum.

#### Beat Recognition versus Rest

The sensorimotor temporal processing task with body movement recruited the auditory dorsal pathway and vestibular cortex, as predicted in the contrast Beat Recognition > Rest, whole-brain corrected at *p*_(FWE)_ < 0.01 (**Table 3, Figure 3**, right panel). The key active regions for auditory-motor integration included: bilateral primary and belt auditory cortex (BA41/42), left posterior superior temporal gyrus (pSTG, BA22), bilateral inferior parietal lobule (IPL, BA40), bilateral premotor cortex and right supplementary motor area (BA6). Two regions of dorsolateral prefrontal cortex (DLPFC) were also activated: bilateral inferior frontal gyrus (IFG, BA9/45), and bilateral middle frontal gyrus (MFG, BA46). Subcortical regions that are integral to the auditory-motor timing circuit were activated, including: bilateral basal ganglia (caudate, claustrum), left anterior and bilateral posterior cerebellum, and bilateral thalamus (see **Table 3** for details). Importantly for the goals of the present study, the left parieto-insular vestibular cortex (PIVC, BA13) was significantly activated in SC and EB groups during Beat Recognition (region of interest analysis to follow).

**Table 3.**
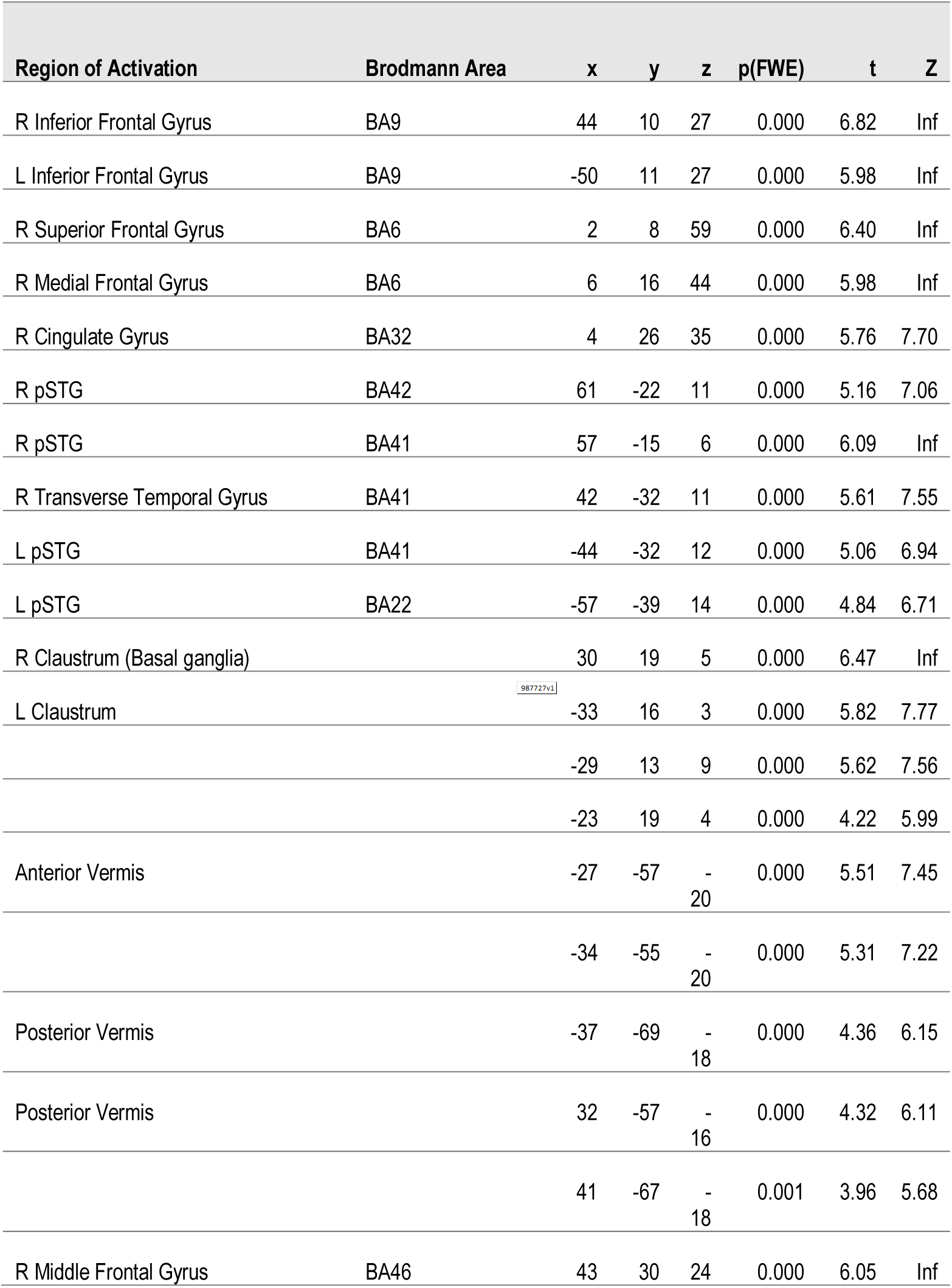
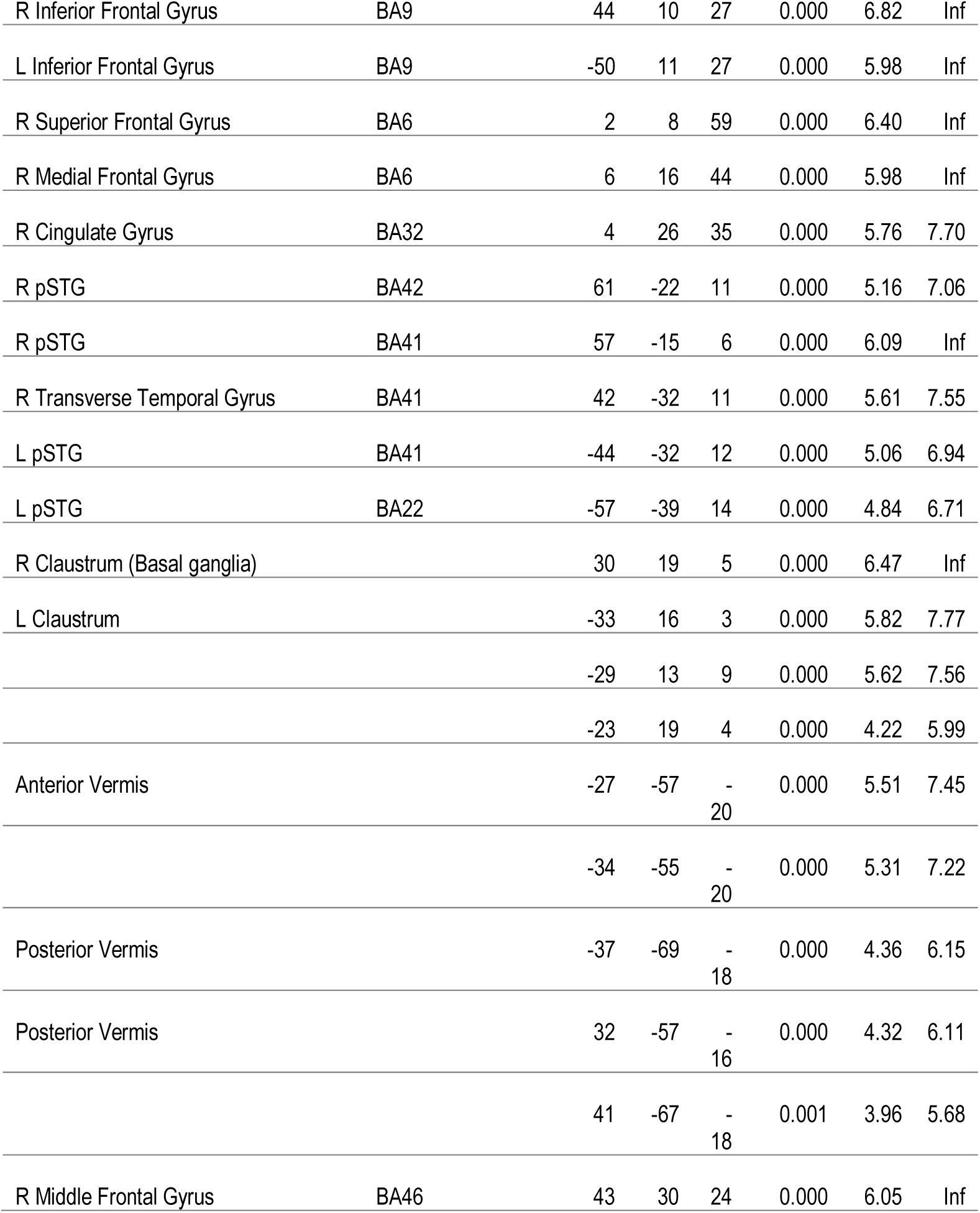
Talairach coordinates for brain regions active in Beat Recognition > Rest, whole-brain corrected at *p(FWE)* < .01.

#### Beat Recognition versus Tapping

To examine the regions of significant activation that were specific to the task of Beat Recognition beyond those activations elicited by listening and producing a motor response in the Tapping task, we performed the contrast Beat Recognition > Tapping, whole-brain corrected at *p*_(FWE)_ < 0.05 **(Table 4a)**. This resulted in activation of: bilateral middle frontal gyrus (BA9/46), bilateral inferior frontal gyrus (BA9), bilateral medial superior frontal gyrus (BA8), and right anterior insula (BA13; **Figure 4**, left panel). Finally, the contrast Tapping > Beat Recognition, whole-brain corrected at *p*_(FWE)_ < 0.05 resulted in significant activation of the anterior and posterior vermis, an unpaired portion of the cerebellum associated with bodily posture and locomotion **(Table 4b; Figure 4**, left panel).

**Table 4.**
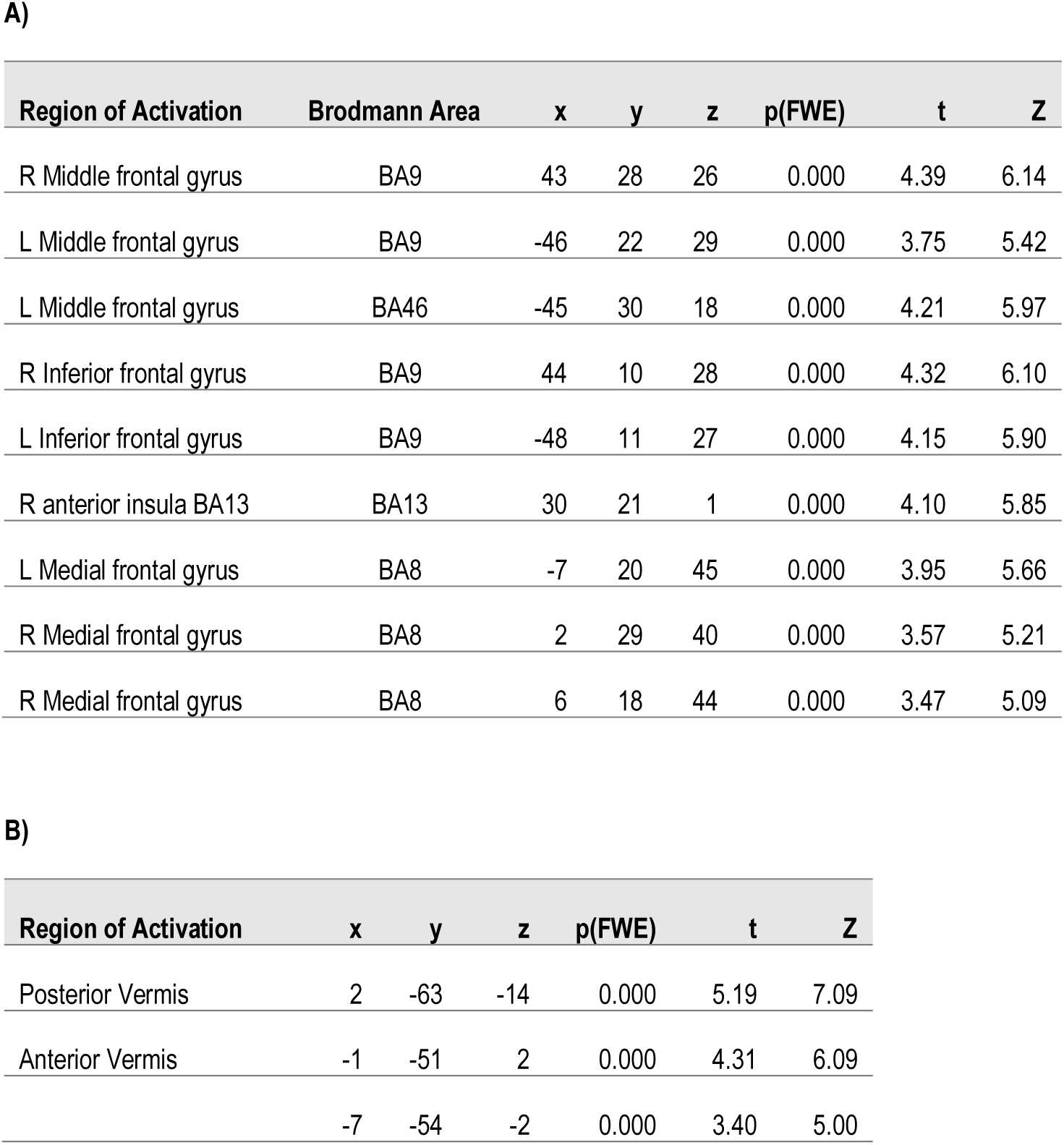
Talairach coordinates for Beat Recognition versus Tap. **A)** Talairach coordinates for brain regions significantly active in the contrast Beat Recognition > Tap, whole-brain corrected at *p(FWE)* < .05. **B)** Talairach coordinates for brain regions significantly active in the contrast Tap > Beat Recognition, whole-brain corrected at *p(FWE)* < .05.

**Figure 4.**
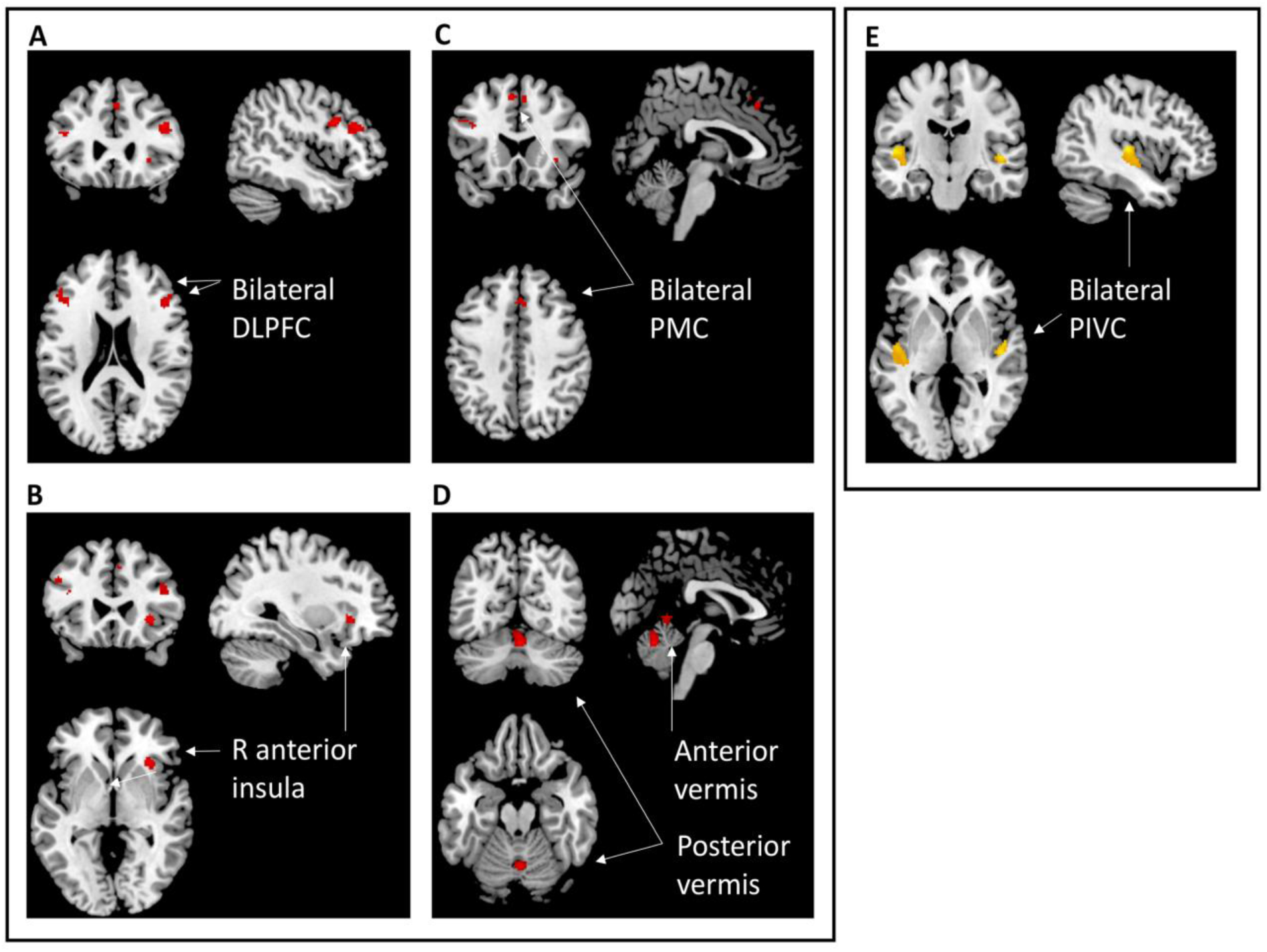
**(A-D)** Beat versus Tap in EB and SC. The contrast Beat > Tap, whole-brain corrected, *p*_(FWE)_ < 0.05 activated bilateral DLPFC (BA9/46), bilateral PMC (BA8), and right anterior insula (BA13). The contrast Tap > Beat, whole-brain corrected, *p*_(FWE)_ < 0.05, activated anterior and posterior vermis. **(E)** ROI analysis of bilateral PIVC (defined by coordinates from Kirsch et al. 2016) for the contrast Beat Recognition versus Rest, in EB and SC. PIVC showed significant bilateral activation, *p*_(FWE)_ < 0.05 in SC; this activation was not significantly different in EB.

#### Vestibular Cortex

To identify the contribution of vestibular cortex, we performed a region of interest (ROI) analysis on bilateral PIVC using a 5 mm^3^ spherical ROI with the center coordinates on the left (x = −42, y = −16, z = 0) and right (x = 44, y = −10, z = 6) as defined by Kirsch and colleagues (2016). According to these authors, the PIVC is composed of posterior ventral insula (Ig1) and retroinsular parietal operculum, which were determined to have functional connectivity to seed regions in bilateral vestibular nuclei (Kirsch et al. 2016). PIVC has been shown to respond to vestibular stimulation in interaction with other sensory modalities (visual and somatosensory; Grüsser et al., 1990). Here, the PIVC showed significant activation for the contrast Beat Recognition versus Rest at threshold *p*_(FWE)_ < 0.05 in SC and EB (**Figure 4**, right panel), and the activation was not significantly different between SC and EB groups.

#### Correlations Between Cortical Activation and Behavioral Measures

We performed exploratory correlational analyses between brain activation patterns during Beat Recognition (versus Rest) and the behavioral measures of Beat Recognition, as well as Physical Spatial Activity. Across EB and SC participants, the degree of activation of PIVC showed a positive correlation with performance on the Beat Recognition task (*r* = 0.48, *p* < 0.007; **Figure 5a**), as did activation of IPL (*r* = 0.48), PMC/SMA (*r* = 0.55), and putamen (*r* = 0.57). Most of these correlations of brain activation levels with the behavioral measure of Beat Recognition were somewhat stronger in the EB group (PIVC, *r* = 0.5; IPL, *r* = 0.48; premotor/SMA, *r* = 0.61; putamen, *r* = 0.68), as compared to the SC group.

**Figure 5.**
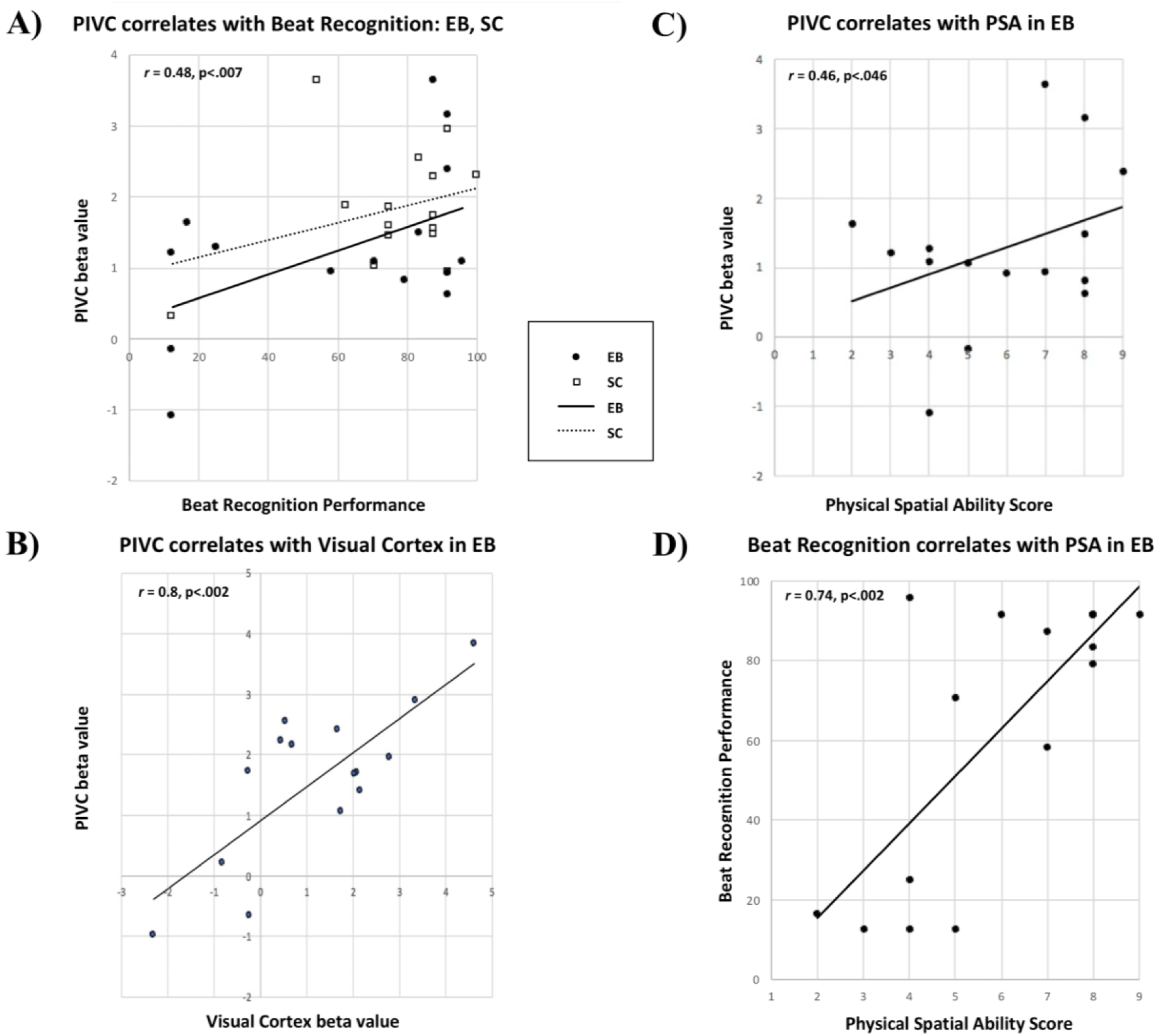
Correlations between fMRI data and behavioral measures. **(A)** In both EB and SC groups, bilateral PIVC activation correlated with performance on the Beat Recognition task. **(B)** In the EB group, activation of PIVC correlated with activation of visual cortex (BA19). **(C)** In the EB, activation of PIVC correlated moderately with physical spatial activity (PSA) score. **(D)** In EB, Beat Recognition performance correlated with PSA score.

In EB specifically, activation of PIVC correlated strongly with activation of visual cortex BA19 (*r* = 0.8, *p* < 0.002; **Figure 5b**). Activation of PIVC correlated marginally with the physical spatial activity score (*r* = 0.46, *p* < 0.046; **Figure 5c**). Finally, the EB group showed a strong correlation between the measure of physical spatial activity (as described in the section on Participants; see **Table 1**) and Beat Recognition performance (*r* = 0.74, *p* < 0.002; **Figure 5d**).

## Discussion

The recruitment of the visual cortex in the early blind for the task of beat recognition reflects crossmodal plasticity in the context of musical rhythm perception. The present study complements findings that the early blind show visual cortex activation for tasks of auditory spatial processing (Rauschecker, 1995; Bushara et al., 1999; Weeks et al., 2000; Gougoux et al. 2005; Collignon et al. 2009; Dormal et al. 2012; Renier et al. 2014), however it is the first study to our knowledge to show crossmodal plasticity for sensorimotor temporal processing. Of particular importance in the processing of temporal sequences during head or body movement is the strong relationship between the degree of activation of visual and parietoinsular vestibular cortex in the early blind.

Despite the close coupling of the visual and vestibular systems, little attention has been paid to the effects of early visual deprivation on vestibular function or on the integration of vestibular input with spatiotemporal auditory and motor information. Yet the available evidence has provided support for the importance of physical spatial activity (Seemungal et al. 2007) and mobility training in the early blind, highlighting the potential for remediation due to cortical plasticity (Rauschecker, 1995). Use versus ‘immobilization’ is a key determinant in cortical plasticity, as seen with tactile perception in the blind (Sathian, 2000; Lissek et al. 2009). Another view posits a role for the visual system as a calibrator for auditory and spatial information, and this view is supported by findings that early visual deprivation can impair spatial navigation, proprioception, and sound localization (Knudsen et al. 1991; Gori et al. 2014a; Cappagli et al. 2017). However even in the latter view it has been suggested that the tactile modality can be utilized to recalibrate auditory space, as supported by findings that sensorimotor pointing (with auditory-tactile feedback) improves sound localization (Gori et al. 2014b). Given the evidence cited above of the importance of movement and utilization in rehabilitation of the blind, there seems to be a great potential for the role of vestibular and motor feedback in the calibration of auditory space and time.

The results of the present study are consistent with the idea that visual experience is not a requisite for sensorimotor spatiotemporal processing, as in rhythmic entrainment. The common activation of the ADS timing network between early blind and sighted control groups suggests that sensorimotor integration for beat recognition is primarily an ADS function, and that it is largely preserved after early visual deprivation. The cortical dorsal auditory pathway areas (pSTG, IPL, PMC, and DLPFC) and their subcortical associated networks (basal ganglia, cerebellum) which were found to be engaged, reflect the timing, anticipation, sensorimotor integration and motor planning components of the task (Rauschecker and Scott, 2009; Rauschecker, 2011).

The fact that rhythmic entrainment originating from body movement also activated PIVC is a significant addition to our understanding of sensorimotor integration and control in the ADS. Vestibular input could make an important contribution to spatiotemporal processing in the visual and auditory ‘where’/’when’ stream. Relayed via the vestibular nuclei of the cerebellum and its vermis (as part of the vestibular cerebellum), this information may in fact be crucial for setting up spatial forward models as postulated for implementation in the dorsal stream (Rauschecker and Scott, 2009; Rauschecker, 2018). The finding that the ADS extends to vestibular cortex when body and head movement are involved in learning rhythm is consistent with prior behavioral evidence that rhythmic entrainment—or, ‘feeling the beat’—is an auditory-vestibulomotor phenomenon (Phillips-Silver & Trainor, 2008; Trainor et al., 2009).

The present finding also indicates that vision was not required to calibrate vestibular input in this task. However, the extent of involvement of the vestibular cortex was more varied in the blind and correlated strongly with the activation of visual cortex (BA19), as well as with a measure of physical spatial activity. These results together suggest that physical spatial activity—involving vestibular stimulation—might mediate the effects of early visual deprivation on sensorimotor spatiotemporal processing via crossmodal plasticity. That the extent of activation of PIVC correlates with behavioral performance in both EB and SC groups may indicate that the involvement of PIVC in the auditory dorsal stream is typical, and consistent with its sensorimotor and spatiotemporal nature.

Although there was significant activation of basal ganglia in the Beat Recognition task, there was less prominent activation than expected in the putamen specifically. The putamen did not emerge from the contrast Beat Recognition > Rest at the threshold reported above, although the right putamen did emerge as significantly active for the same contrast when whole-brain corrected at a threshold of *p*_(FWE)_ < 0.05. The putamen is known to be central to beat processing tasks, and it has been suggested that its role is particularly modulated by the apparent strength of the perceived beat, and the ability to predict upcoming beats based on an internal model (Grahn and Rowe, 2013). Yet the stimuli in the present study were specifically designed to be bistable (that is, possible to interpret with either binary or ternary strong beats), so that vestibulomotor input during training was required to recognize the beat structure of the rhythms during testing. Thus, it could be that this feature of bistability actually diminished the role of the putamen, relative to other regions responsible for sensorimotor integration.

The comparison between the two entrainment tasks of Tapping and Beat Recognition allowed us to observe which regions of the ADS were specific to motor generation in the context of the ambiguous auditory pattern, and which were specific to the task of recognizing the beat as learned during the auditory-vestibulomotor training. Tapping, more than Beat Recognition, engaged anterior and posterior regions of the cerebellar vermis, an unpaired portion of the cerebellum associated with bodily posture and locomotion, probably in conjunction with the basal ganglia (Bostan et al. 2010). The five main regions of the cerebellum that receive inputs from the vestibular nerve or vestibular nuclei are the nodulus and ventral uvula, the flocculus and ventral paraflocculus, the oculomotor vermis of posterior lobe, vermal lobules I and II of the anterior lobe, and the deep cerebellar nuclei. While the anterior vermis is part of the vestibular cerebellum (Barmack, 2003), the posterior regions reflect prediction and motor execution components of the task (Buckner, 2013; Sokolov et al. 2017). Thus, the recruitment of anterior and posterior vermis during Tapping might suggest subcortical predictive and motor function along with vestibular input.

Beat Recognition, more than Tapping, activated bilateral DLPFC and PMC, which reflect the cognitive, motor planning and monitoring nature of the task of recognizing and indicating the ‘felt beat’ (Rauschecker and Scott, 2009). Beat Recognition also more strongly activated right anterior insula, which is thought to reflect interoception (Critchley et al. 2004; Craig, 2009), and which also emerges in numerous studies on rhythm perception (e.g., Grahn and Rowe, 2013). Also in Beat Recognition more than in Tapping, the right caudate and left IPL were significant at the cluster level, although they did not survive whole-brain correction. While the overall activation of the ADS network was equivalent between EB and sighted groups, the greater strength of the correlations observed in the EB between brain activation levels and behavioral measures of Beat Recognition may point to a form of enhancement due to compensatory plasticity. This possibility warrants further investigation in studies especially on sensorimotor spatiotemporal processing in the early blind.

In light of the effect of full body movement—in particular, vestibular input—on auditory spatiotemporal processing via the ADS, we propose two points for future investigation. The first is to further consider the role of vestibular cortex in temporal processing when the head or body is moving in space—that is, auditory-vestibulomotor temporal processing (see also Todd, 2015). Such activities include rhythmic entrainment in music and dance as prime examples, but also include sports, locomotion, and navigation. Second is to examine the potential for physical spatial training to induce crossmodal changes based on synaptic plasticity, and improve spatiotemporal processing, as well as the perception and utilization of vestibular information in the early blind. The restricted movement that is common in young blind children is likely to cause deficiencies in the development of the vestibular system (and thus in certain spatial functions). Nevertheless, through the practice of physical spatial activities—including music and dance—the vestibular system may be highly adaptable, and critically important for rehabilitation, orientation and mobility training.

## Conclusions

Rhythmic entrainment, when it involves full body movement—as in *‘*feeling the beat’—is an auditory-vestibulomotor phenomenon which engages the ADS for timing and action, with an extension to parietoinsular vestibular cortex. This ability is preserved in the early blind, however its development may depend somewhat on experience with physical spatial activities. The present findings support the idea that vestibular information is integral to sensorimotor processing of temporal information such as in musical rhythm, and highlight the possibility that musical rhythm could provide an opportunity for mobility training and vestibular processing in early blindness.

